# Microfluidic characterization of macromolecular liquid-liquid phase separation

**DOI:** 10.1101/2020.06.16.154518

**Authors:** Anne Bremer, Tanja Mittag, Michael Heymann

## Abstract

Liquid-liquid phase separation plays important roles in the compartmentalization of cells. Developing an understanding of how phase separation is encoded in biomacromolecules requires quantitative mapping of their phase behavior. Given that such experiments require large quantities of the biomolecule of interest, these efforts have been lagging behind the recent breadth of biological insights. Herein, we present a microfluidic phase chip that enables the measurement of saturation concentrations over at least three orders of magnitude for a broad spectrum of biomolecules and solution conditions. The phase chip consists of five units, each made of twenty individual sample chambers to allow the measurement of five sample conditions simultaneously. The analytes are slowly concentrated via evaporation of water, which is replaced by oil, until the sample undergoes phase separation into a dilute and dense phase. We show that the phase chip lowers the required sample quantity by 98% while offering six-fold better statistics in comparison to standard manual experiments that involve centrifugal separation of dilute and dense phase. We further show that the saturation concentrations measured in chip are in agreement with previously reported data for a variety of biomolecules. Concomitantly, time-dependent changes of the dense phase morphology and potential off-pathway processes, including aggregation, can be monitored microscopically. In summary, the phase chip is suited to exploring sequence-to-binodal relationships by enabling the determination of a large number of saturation concentrations at low protein cost.

## Introduction

Liquid-liquid phase separation (LLPS) has been identified as the underlying driving force for membraneless compartmentalization in cells ^1, 2^. LLPS plays key roles in fundamental cell biological processes such as cell signaling ^3^, stress responses ^4-6^, transcription ^7^, RNA splicing ^8^, RNA metabolism ^9^ and clustering of receptors at cell membranes ^3, 10^. Importantly, recent findings indicate that phase separation and its dysregulation can be linked to cancer ^11, 12^, neurodegenerative diseases ^13, 14^ and aging ^15^.

Phase separation is mediated by multivalent interactions, which enable the formation of a non-covalently networked, dense phase (with concentration *c*_d_) that coexists with a dilute phase (with concentration *c*_sat_, also called the saturation concentration). *c*_d_ and *c*_sat_ are often modulated by temperature, pH, solutes and binding partners, generating solution condition-responsive phase behavior, which can be mapped as two-dimensional phase diagrams.

Disease-associated mutations in phase-separating proteins can shift the saturation concentration (*c*_sat_), i.e., the threshold concentration beyond which the protein forms a dense phase. If protein function depends on its colocalization with partners in a biomolecular condensate or on its segregation from partners through selective phase separation, then an alteration of the saturation concentration can be detrimental and can lead to loss or gain of function, respectively. Further, fusion of phase separating protein fragments to the DNA binding domains of transcription factors, as a result of chromosomal translocations ^12, 16^, can result in *de novo* gain of phase behavior and aberrant transcription. This mechanism is thought to be the cancer-initiating step in cells with FET family fusions ^17, 18^. To understand the impact of disease mutations and fusion events, and to improve our understanding of the involvement of phase separation in physiological processes, there is a need to characterize the biophysical nature of the interactions that drive LLPS, to extract the pairwise interaction strengths of these interactions and to build models that quantitatively predict phase behavior.

Our ability to predict phase behavior, i.e. to predict the dilute and dense phase concentrations as a function of temperature, salt concentration, pH or binding partners, is in its infancy. Notable efforts have included the systematic determination of saturation concentrations of FUS and FET family proteins and of designed sequence variants. The data was used to derive an analytical expression of how the saturation concentration depends on the number of tyrosines and arginines in the sequences, assuming that they are the major adhesive elements ^19^. Other efforts include the development of models for electrostatically mediated phase separation in polyampholytes, i.e. of sequences that contain high fractions of negatively and positively charged residues, as a function of how these oppositely charged residues are patterned in the sequence ^20-22^. Full temperature-dependent binodals were predicted in this case. In recent work, we determined the major stickers in the intrinsically disordered prion-like domain of hnRNPA1, determined their microscopic interaction strengths and developed a numerical stickers-and-spacers model that is predictive of full coexistence curves (i.e. binodals) for this flavor of IDR ^23^. Overcoming the current barriers to accurate prediction of phase separation of IDRs of all flavors will require the development and refinement of general theoretical models, and their parameterization will require the precise determination of binodals for a large number of proteins and sequence variants. The ability to predict phase behavior will also enable the development of phase-separating proteins with desired material properties and solution-responsiveness.

Constructing phase diagrams is challenging, mainly because large quantities of the biomolecule of interest is needed, and because of the time requirement for repetitive, manual measurements. Hence, the determination of saturation concentrations has been largely limited to single conditions ^19^ and full binodals have been determined only for a handful of proteins ^23, 24^. Most binodals are constructed from a combination of turbidity assays that are used to determine cloud points ^19, 23, 25-27^ and centrifugal separation of the dilute and dense phases coupled with the spectroscopic determination of their protein concentration ^19, 23, 28, 29^. Disadvantages of these techniques include (i) the requirement for large material quantities, (ii) the difficulty to identify potential off-pathway processes (e.g. aggregation) and (iii) a lack of information on differences in droplet morphology between different conditions, which may point to interesting physicochemical effects. An alternative approach that allows visualization of the dense phase is the preparation of a dilution series of protein samples, each of which are scored for the presence of two phases by microscopy ^30-32^. This approach, however, is not fully quantitative because discrete protein concentrations are assessed for whether they contain two phases or not, and the saturation concentration is not directly determined. Depending on the choice of concentration grid points, the real saturation concentration may be missed by a large value, in which case the resulting phase diagrams are not suitable for parameterizing quantitative models.

Extensive microfluidic engineering has been applied to study macromolecular phase transition phenomena, ranging from normally open ^33^, to normally closed valves ^34^, as well as emulsion droplet workflows ^35, 36^ for combinatorial composition titrations. Alternatively, semipermeable membrane transport through permeation of water vapor ^37-39^ or dialysis ^40, 41^ has been successfully used to study phase transitions. However, all these approaches were constrained by complex device fabrication, operating equipment requirements, consumption of comparably large quantities of precious biological sample, by low throughput or a combination thereof.

Here, we report a microfluidic device for accurate and precise determination of full binodals over a broad range of conditions. Through capillary valving, we achieve loss-free sample loading such that a total sample volume of less than 10 µL enables mapping of saturation concentrations at five conditions with twenty replicates in a single experiment. All fluidic operations are achieved without microfluidic controls such as actuated valves or pumps, rendering the system suitable for non-expert users and directly usable with routine microscopy infrastructure. We show that we can image phase separation and determine saturation concentrations that vary three orders of magnitude and that the resulting data is in agreement with previously recorded saturation concentrations determined by absorbance and light scattering measurements. In parallel, time-dependent information on potential morphological changes of the dense phase is collected. The chip affords the opportunity to explore sequence-to-binodal relationships at low cost and to monitor the morphology of condensed states, bringing us closer to our goal of generating experiment-grounded theories of protein phase separation.

## Results

### Microfluidic chip design and fabrication

The PDMS-glass chip compartmentalizes biomolecular phase separation mixtures into nanoliter sized emulsion wells and is actuated by hydrostatic pressure-driven flow (Fig. 1). It consists of five units, each made of twenty individual sample chambers that are connected by a bypass to allow the measurement of five sample conditions simultaneously. This enables the precise control of solution conditions in space and time while monitoring phase separation in 100 wells in parallel.

**Fig. 1.**
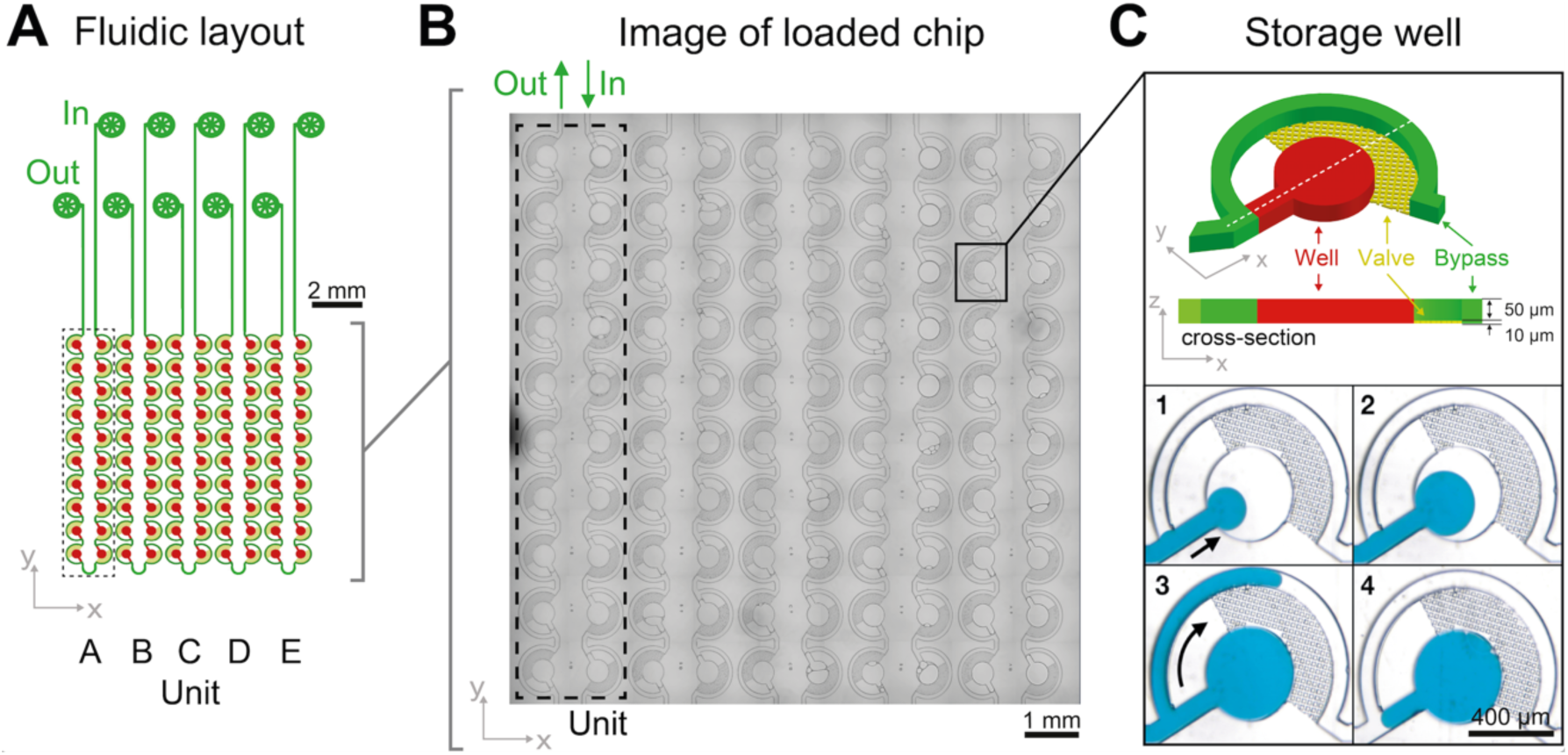
Chip design and loss-free loading principle through capillary valving. (**A**) The LLPS chip consists of 5 units, each of which is comprised of 20 individual sample wells, which are connected by a bypass. (**B**) Micrograph of a loaded chip. (**C**) A zoom of an individual sample well demonstrating the capillary valve loading procedure: After priming with fluorinated oil, the hydrodynamic resistance of the bypass channel (green) diverts sample flow (blue) into the storage well (red) (C1,2). Oil can flow through the capillary valves (yellow), but the aqueous sample cannot. It is instead diverted through the bypass after the well (red) has filled (C3, arrow). (C4) In a final step additional oil is flown through the device to cut each sample well to a defined 8.5 nl volume, set by the 400 µm diameter and 60 µm height of the wells. Detailed design schematics of the fabrication procedures are provided as ESI and in Fig. S1.

Sample is introduced manually through an injection loop for each concentration unit. Chip units are comprised of 20 serially connected store-then-create wells ^38^ (SI design file) in a multi-height configuration ^42^, with capillary valves of 10 μm height, wells and bypass channels of 60 μm height (Fig. S1), and a nominal volume of 8.5 nl per well. All channel walls are treated with amorphous cytop fluoropolymer, which in combination with HFE 7500 oil and PFPE-PEG triblock surfactant has been shown to exhibit long term biocompatibility ^43^. Compared to previous implementations, we introduce a cascaded capillary valve configuration that enables robust manual loading (Fig. 2). The phase chip is a single use device to make it user-friendly and accessible to a broad audience.

**Fig. 2.**
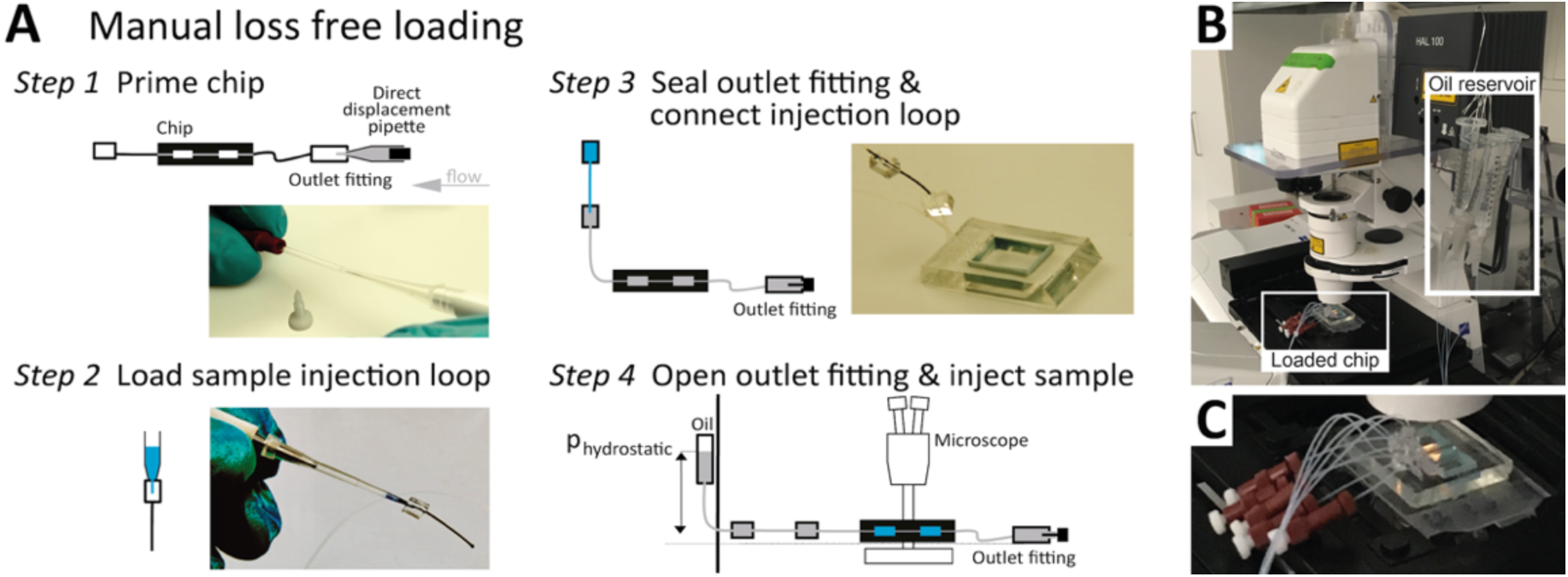
The phase chip enables easy, loss-free loading. (**A**) Steps to manual, loss-free loading. Step 1: Insert about 4 cm of 0.3/0.75 mm ID/OD PTFE tubing into all channel outlets. Terminate each outlet tubing with a P-881 fitting and a dead-end plug. Prepare inlet tube connectors using an identical tubing segment capped with a PDMS cube of 5 mm edge length with a 0.75 mm punch-through hole. Prime all channels and connect inlet and outlet tubing by flushing 25 µl droplet oil, using a 25 µl direct displacement pipette. Note, removing all bubbles from the fluidic circuit is best done by flushing from the outlet. Seal the outlet by inserting a P-881 dead-end plug. Step 2: Aspirate the sample into a 2 µl manual air displacement pipette. Connect pipette inlet to the injection loop comprising of 2 cm, 0.3/0.75 mm ID/OD PTFE tubing capped with a PMDS connector block and inject the sample. Step 3: Eject the pipette tip and connect the injection loop pipette assembly to the inlet tube. Step 4: Remove the tip and connect a HFE-7500 hydrostatic pressure reservoir to the injection loop. (**B**) After positioning the chip on the microscope and defining the microscope image acquisition, (**C**) open the P-881 dead-end plug slightly to allow for the hydrostatic pressure-driven flow to load the wells. After all wells are filled close the dead-end plugs again.

### Measurement principle of evaporation-based phase behavior quantification

Next, we tested whether the saturation concentration of biomolecules can be reliably determined with this phase chip implementation. We are taking advantage of the simple fact that water evaporates through the semipermeable PDMS device wall over time ^37, 38, 42^ resulting in solution volume decrease in the storage well and concomitant increase of protein and buffer solute concentrations (Fig. 3). Evaporated water is replaced by hydrostatic oil from a reservoir to prevent unwanted side effects of protein surface/air interactions. Upon reaching the saturation concentration the sample undergoes phase separation and demixes into dilute and dense phases. The height of the sample volume remains constant during evaporation while the radius of the sample volume decreases. Each well has “pancake”-like dimensions with a diameter of 400 μm and height of 60 μm. Surface tension minimizes the interfacial area and an emulsion droplet with a diameter exceeding the well height is distorted into a cylindrical shape. If a cylindrical droplet with a radius (*r*) equal to the well height (*h*) is considered as the lower boundary of a droplet volume that would not detach from the bottom and top of the well (*V*_*min*_), then the dynamic range of cylindrical volume changes with constant height approximation, is given as

**Fig. 3.**
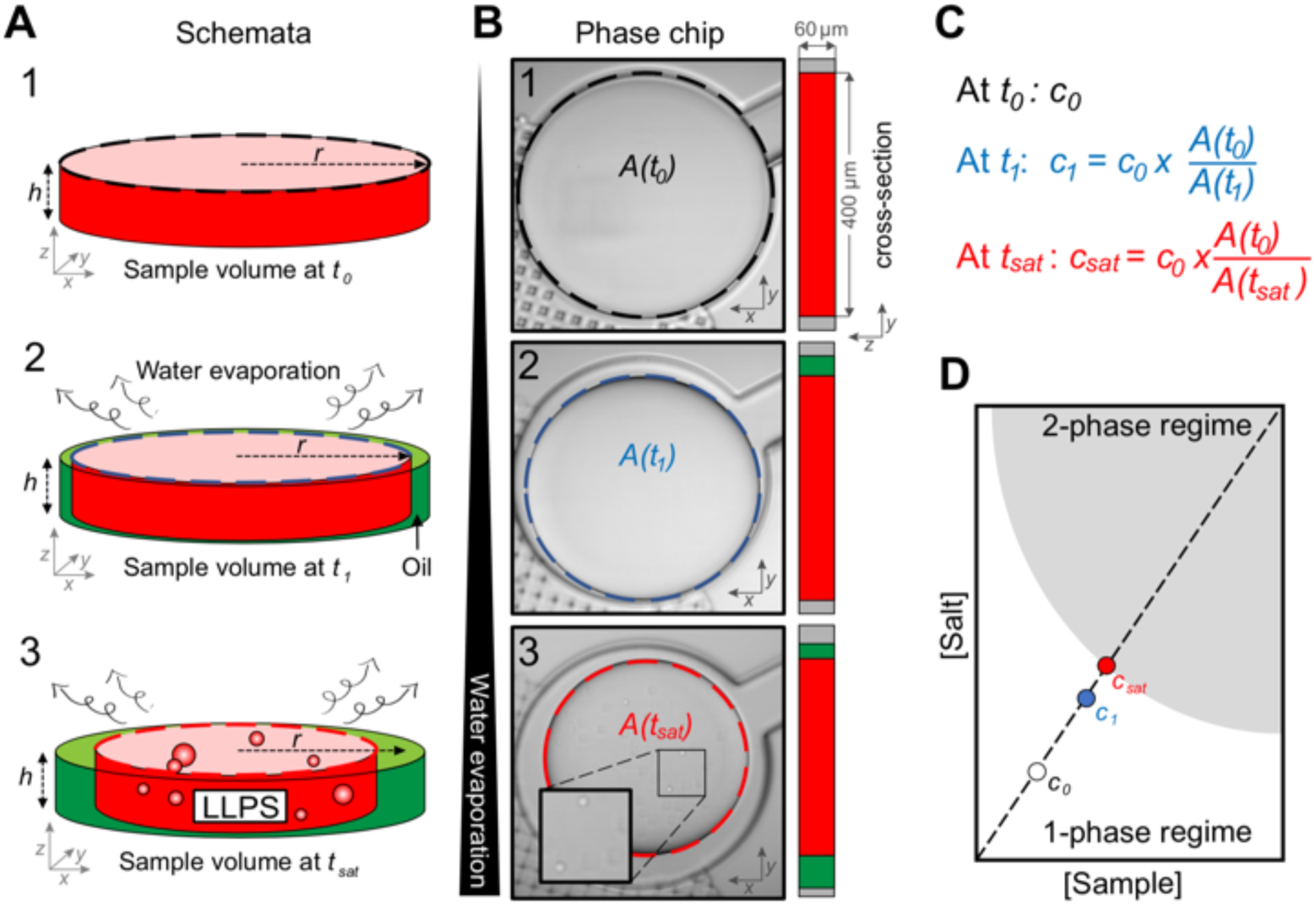
Determination of saturation concentration (c_sat_) from sample volume. (**A**) At the beginning of the experiment, the sample fills the entire chamber. Water evaporates over time at constant temperature and the sample volume inside the chamber shrinks while the overall sample concentration increases (step 2). The height of the sample droplet is maintained by the channel geometry (no detachment from top), hence only the radius changes. Evaporating water volume is replenished by oil. When the sample concentration reaches c_sat_, the solution phase separates (step 3). (**B**) Live images at time points corresponding to (A) The shrinking surface area of the sample A(t) is determined by image analysis. (**C**) Calculation of the saturation concentration c_sat_ based on the starting concentration of the sample (c_0_) at t_0_ and the starting and current sample surface areas. (**D**) Linear dependency of the sample concentration and salt concentration for any given time point based on the equation in (C).

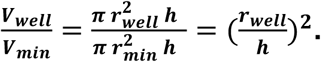

We selected this ratio to be about 10fold, while experimentally never exceeding a five-fold concentration increase. E.g. if a starting concentration of 10 μM is loaded, the protein could have a saturation concentration of up to 100 μM and it could still be accurately measured. Given that most physiologically phase-separating proteins have much lower saturation concentrations and given further that the loading concentration can be increased, we conclude that this is sufficient for a wide range of experimental targets. At the same time, reducing the well height and/or increasing the radius would increase the dynamic range of quantifiable volumes, such as realized in the 20nl-chip (600 μm diameter, 50 μm high wells) of Selimovic et al. which allows a 36fold decrease in volume ^38^. The sample concentration at any given time can thus be inferred from the change in radius of the aqueous compartment. The concentration at the onset of phase separation is the saturation concentration, *c*_sat_, and is calculated by multiplying the sample concentration at time point zero *c*0 with the ratio of sample areas at *t* = 0 and at the time of phase separation,

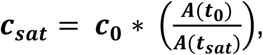

to ultimately construct a phase diagram (Fig. 3C, D).

Since water vapor permeation is linearly proportional to distance, we reduced the PDMS ceiling thickness to about 600 µm to obtain an optimal evaporation rate to reach half the initial volume after ∼7.5 hr (Fig. S2). A thin chip offers the advantage of faster evaporation and phase separation earlier in the experiment, while a chip that is too thin becomes more delicate to produce and is more prone to compliant deformations and rupture of the device ceiling. Another downside of a fast evaporation rate is its limiting effect on the precision of the microscopic acquisition, as the area of observed positions and/or the temporal resolution must be reduced with enhanced sample evaporation. Alternatively, water may be extracted through continuous oil flow as has recently been demonstrated for monitoring phase separation with a microfluidic device ^39^.

We then tested whether a protein sample that was passed through the chip experienced sample loss due to protein adsorption on tubing or device surfaces. Any non-specific binding would compromise the precision of the determined saturation concentration given its dependence on the initial concentration. We determined sample concentrations before and after passing the sample through the chip and the concentrations were identical. We highly encourage all users to repeat this control for their biomolecule of interest to ensure a loss free sample loading. Also, no fluorescently labeled protein was observed to absorb to the device surface (Fig. S3).

### Accurate microfluidic determination of binodals

To test whether the microfluidic chip can be used to determine full binodals accurately and reproducibly, we used the PEG/ammonium sulfate system whose phase behavior has been extensively characterized ^37, 44^. A broad range of pairwise PEG and ammonium sulfate concentrations were loaded and incubated on chip and their pairwise saturation concentrations were determined (Fig. 4) from three to four independent experiments. Two microscope objectives with different magnifications (10x and 20x) were used for comparison. A manual combinatorial dilution series (Fig. 4A) was preferred over a microfluidic formulator ^45, 46^. Macromolecular solutions at concentrations when they are prone to undergo LLPS diverge vastly in composition and viscosity, rendering it difficult to achieve a universal fluidic design that is compatible with all possible solution conditions and also suitable for manual operation by users without prior microfluidics expertise. All saturation concentrations obtained from observations of individual microfluidic chambers are shown in Fig. S4. The average values and their associated errors determined on chip agreed well with previously reported turbidity cloud point measurements ^44^. *c*_sat_ values obtained using the objective with 10x vs. 20x magnification were identical within error, showing that the lower magnification objective is sufficient for the precise determination of the onset of phase separation.

**Fig. 4.**
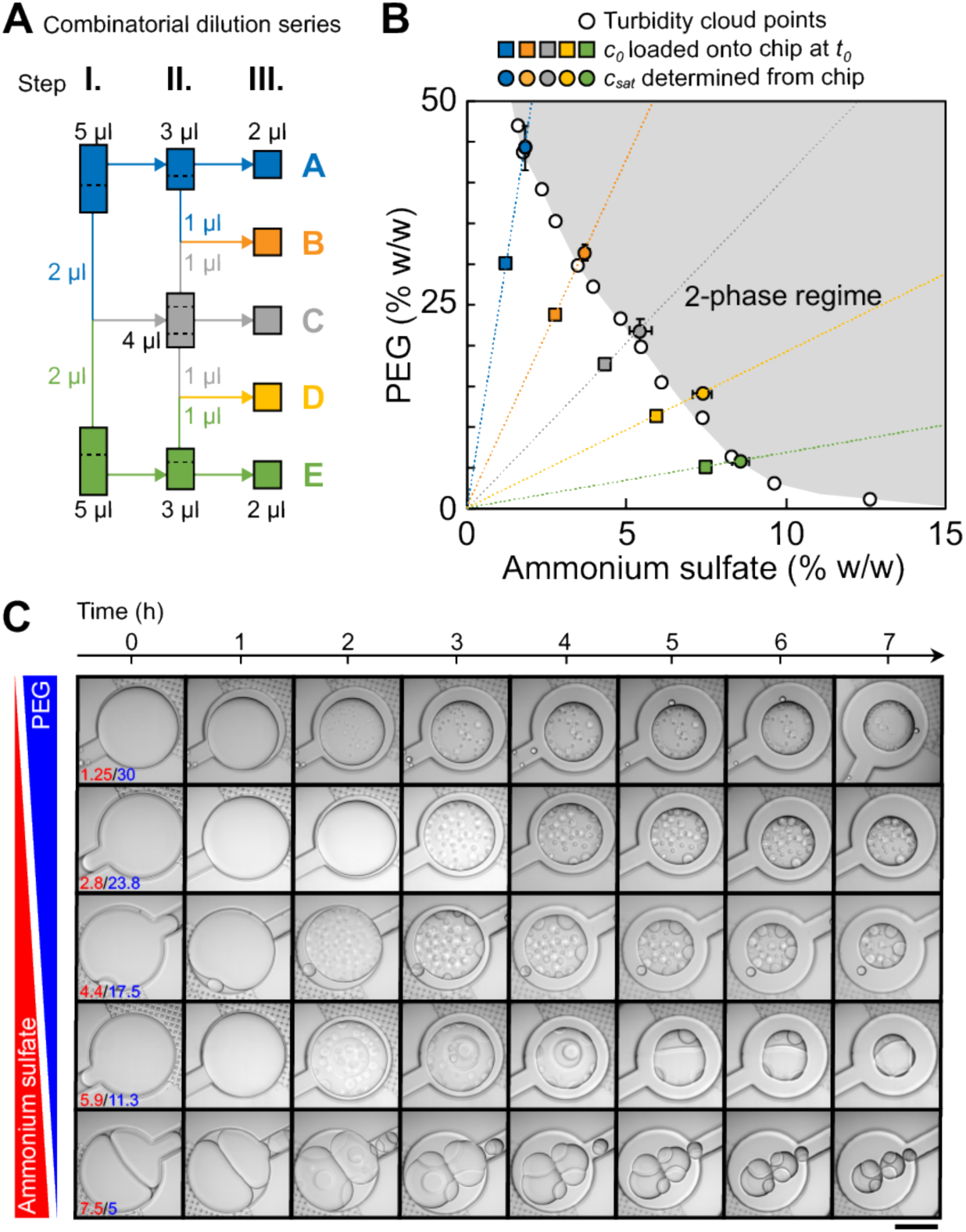
Microfluidic determination of saturation concentrations of pairwise PEG/ammonium sulfate (AS) combinations. (**A**) Schematic of the three-stage combinatorial dilution series used to yield the five compositions from the two initial conditions. A volume of 1 µl was considered as the lower limit for accurate manual pipetting. The dotted lines indicate how the volume was split. (**B**) c_sat_ determined on chip were compared to previously reported data points (empty circles) measured via turbidity cloud point assays ^44^. Transitions from 10 wells from three independent experiments per condition were analyzed. Error bars are the standard deviations of the mean of all transitions. Starting concentrations of PEG and ammonium sulfate are shown as squares. A smooth line is drawn through the measured c_sat_ values to guide the eye. (**C**) Time dependent onset of LLPS and morphological changes of pairwise PEG/ammonium sulfate phase separated droplets are shown (Movie S1). Images were taken with LSM800 using a 20x objective. Scalebar represents 200 µm.

A tile scan of the full 1 cm^2^ chip area took three minutes using the 10x objective, and consecutive timepoint scans were taken without waiting periods. Due to continued water evaporation during acquisition of a single tile scan, we determined the difference of the sample volume (and therefore *c*_*sat*_) between the image that contains the first signs of dense phase (which we define as the transition point) and the image one time point earlier (the pre-transition point). The difference was less than 3% for all five conditions (Fig. S5), thus a minimal error of below 3% was introduced by the averaging nature of the microscopic measurement modalities.

### Observation of dense phase morphologies

The left arm of binodals is typically reconstructed from turbidity assays ^19, 25^ or by separating dilute and dense phases by centrifugation ^6, 19, 28^; possible morphological changes of the dense phase over time are not accessible with these approaches. The phase chip does not only provide access to precise saturation concentrations but also time-dependent information on the morphology of the dense phase over the time course of the measurement. We observed changes in morphology of several PEG/ammonium sulfate dense phases at specific conditions (Fig. 4C, e.g. PEG/AS 5/7.5 (% w/w) and 11.25/5.9 (% w/w)). Many individual dense phase droplets gave way to few large dense phase droplets, indicating their rapid fusion. After 3 hours, light and dense phase of the PEG/AS 11.25/5.9 (% w/w) condition clustered in a way to limit common interfaces, indicating a high surface tension; we observed a similar behavior for PEG/AS 5/7.5, but less pronounced. Other pairwise PEG/AS concentrations retained the morphology immediately observed after initial phase separation.

Our data shows that the microfluidic chip enables the reliable determination of saturation concentrations over a broad range of PEG/ammonium sulfate mixtures and simultaneously provides time-dependent data on dense phase morphology.

### Microfluidic determination of a broad range of saturation concentrations

Next, we tested the performance of the chip on several protein targets that undergo LLPS with different saturation concentrations. We used the well-characterized low-complexity domain of the stress granule-associated RNA-binding protein hnRNPA1 (A1-LCD), which is sufficient for mediating LLPS in vitro ^6^. Binodals of A1-LCD have been characterized by a multi-pronged approach using Fluorescence Correlation Spectroscopy, centrifugation followed by UV-spectroscopic determination of the dilute and dense phase concentrations and cloud point determination by static light scattering ^23^. The saturation concentration of A1-LCD at near physiological conditions at a NaCl concentration of 150 mM at room temperature is ∼80 µM. Many proteins with physiologically relevant phase behavior have saturation concentrations between 1-100 µM ^19^. To test the applicability of the method to proteins with a broad range of saturation concentrations, we included two additional proteins in our analysis, a sequence variant of the A1-LCD with a higher content of charged residues termed A1-LCD^+12D+7R^ and BSA; they have saturation concentrations at 150 mM NaCl and room temperature of 6 µM and ∼3200 µM, respectively. All proteins were purified to homogeneity (Fig. S6). We directly compared saturation concentrations obtained by on chip measurements with values determined by centrifugation to separate dilute and dense phases followed by UV spectroscopic determination of the dilute phase concentration. The A1-LCD and A1-LCD^+12D+7R^ protein stock solutions were kept in buffer without excess salt and were therefore in the one-phase regime. In the test tube, phase separation was induced by either adding NaCl or pairwise PEG/ammonium sulfate concentrations to the protein stock solution. The sample was incubated for 20 min at room temperature and dilute and dense protein phases were separated via centrifugation. The dilute phase was removed and *c*_*sat*_ was determined via UV absorbance at 280nm. The remaining dilute phase was diluted two-fold and 2 μL were loaded onto the chip (Fig. 5A). The chambers were imaged over ∼5 hours to determine the onset of phase separation ^47^. Saturation concentrations determined on chip agreed well with those determined manually in the tube for all three proteins (Fig. 5B-E).

**Fig. 5.**
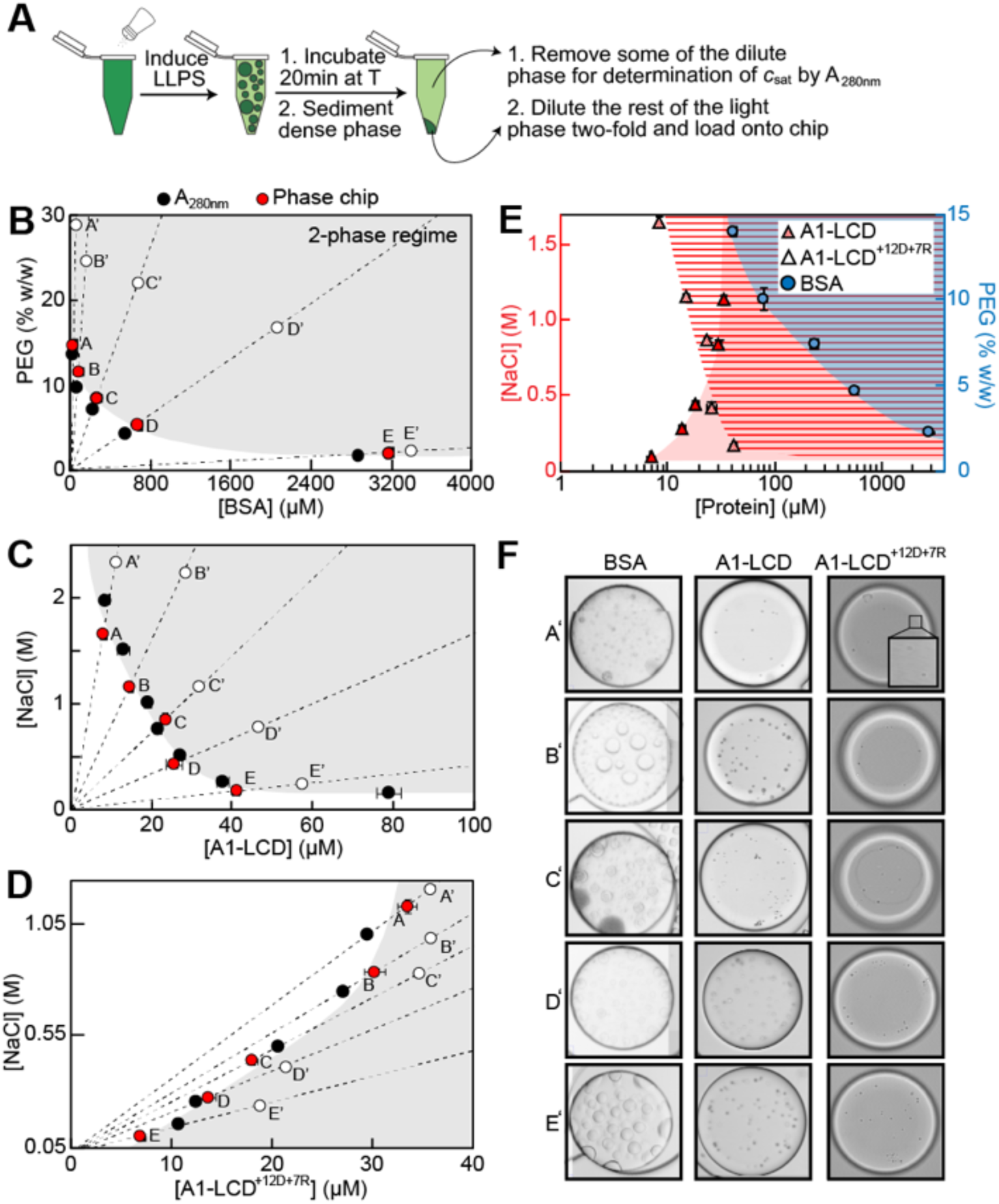
Measurement of saturation concentrations of three biomolecules that undergo LLPS ranging from 6 μM to ∼3200 μM. (**A**) To generate samples, phase separation was induced, and samples were incubated for 20 min at 20 °C followed by centrifugation. For samples containing protein, a part of the dilute phase was removed for UV spectroscopic determination of c_sat_. 2 μL of light phase was diluted two-fold and loaded onto the chip. Saturation concentrations from microfluidic chip measurements were compared to those from bulk assay absorbance measurements at 280 nm (black dots) for (**B**) PEG:BSA, (**C**) A1-LCD, and (**D**) A1-LCD^+12D+7R^. Error bars are the standard deviations of the mean of all droplets used for the analysis. Open circles indicate conditions shown in (F). (**E**) Same data from (B-D) combined into a semi-logarithmic plot, illustrating the large dynamic range spanning three orders of magnitude of the microfluidic phase diagram analysis. Shaded regions are the two-phase regimes of the different proteins. (**F**) Microscopy images of the phase separated proteins taken at the indicated time points (A’-E’, open circles in B-D). Movie S2-4.

All three proteins formed spherical condensates that fused over time, confirming that we indeed characterized the saturation concentration of a LLPS process (Fig. 5F). For some dense phase droplets of the A1-LCD we observed the formation of fibril-like structures over time (Fig. S7). A hexapeptide of the A1-LCD construct was previously shown to act as a steric zipper that seeds A1-LCD fibrillization ^47^ and this is enhanced by phase separation ^6^. Our observation was thus in agreement with our expectations. In conclusion, the microfluidic chip method is applicable to accurate and precise measurements of a broad range of saturation concentrations at minimal protein cost, almost 98% lower than the manual method in test tubes requires.

## Conclusions

We have designed and engineered a microfluidic chip for the reliable determination of saturation concentrations of proteins and other macromolecules. The measured saturation concentrations spanned almost three orders of magnitude. The chip further offers the ability to determine saturation concentrations at several different conditions simultaneously but consumes only small sample quantities. Every condition is assessed twenty times in individual chambers, enabling valuable statistical analyses. Given that the onset of phase separation is determined microscopically, off-pathway or maturation processes can also be monitored and distinguished from LLPS. The chip is broadly applicable and can be operated without dedicated microfluidic control equipment. It uses routine fabrication techniques and after sourcing a wafer it can be fabricated in all labs that have access to an oxygen plasma bonder. This low entry barrier to fabrication and operation combined with the comparably high-throughput measurements of saturation concentrations for many different protein samples renders this device suitable for adaptation in labs without prior microfluidics expertise and will enable the determination of sequence-to-binodal relationships.

A total of 2 µL of sample volume (here 1 µL sample at the dilute phase concentration, two-fold diluted in buffer) is sufficient to load all 20 individual chambers per unit. In our standard manual experiments in tubes, 40 µL of sample with at least a 2.5-fold higher concentration is required for triplicate measurements. Therefore, only one fiftieth of the protein is required for on chip measurements, which equals protein savings of 98% at six-fold better statistics.

Limitations of the microfluidic chip include a tradeoff between image quality and duration of the experiment. Given that the whole chip needs to be imaged at every time point, higher resolution acquisition reduces the number of wells that can be observed simultaneously, which in turn decreases the precision of the determined saturation concentration. This may be counteracted by slowing the evaporation rate in thicker chips. With the automated stage used in this study, a scan of the whole chip, i.e. of 10 x 10 tiles, using a 10x objective required 3-minutes resulting in a 3% error in the determined saturation concentration values. Aggregation-prone proteins present a challenge as they need to remain stable over the course of the measurement, typically several hours. The precision of the determined saturation concentration is furthermore dependent on the concentration ratio of the dilute and dense phase concentrations. If this ratio is larger, i.e. if a protein undergoes phase separation at a relatively low concentration but forms a dense phase with a high density, then the volume fraction of the dense phase will be small and the onset of droplet formation more difficult to determine.

Several other groups have recently used microfluidics approaches to map biomolecular phase diagrams. Knowles and coworkers adopted differential encapsulation to formulate phase separating emulsion droplets that span a wide range of solution conditions ^48^. The droplets were collected and analyzed by epifluorescence microscopy for being in the one-phase or two-phase regime. In this approach, coexistence curves are not directly measured but rather inferred by scanning the solution conditions that gave rise to LLPS. Chilkoti, Lopez and coworkers used water-in-oil emulsion droplets as simple cell-like compartments to probe phase behavior of elastin-like peptides in response to temperature changes to cross and pinpoint the coexistence curve ^49^. Arosio and co-workers extracted water content from phase separating emulsion droplets through continuous oil flow ^39^. Microfluidic mixing techniques have also been used to quickly generate and characterize the phase behavior and permeability of dense phases that quickly mature into solid states ^50^ Microfluidic approaches for mapping phase diagrams have recently been reviewed in detail ^51^.

Future developments of the herein reported phase chip and other devices will include automated image analysis and temperature-control to access phase separation for all physiological temperature ranges ^38^. Further effort will go into the characterization of protein dense phase concentrations on chip through the determination of dense phase volume fractions, dynamic light scattering ^42^ or fluorescence correlation spectroscopy ^23^. Although measurements of dilute phase concentrations have become standard, dense phase concentrations are currently rarely reported, despite the fact that dense phase concentrations are required to develop complete theories that predict phase separation based on the sequence or protein architecture. Furthermore, the microfluidic chip can also be used to study fluorescently labeled macromolecules enabling the characterization not only of homotypic but also heterotypic phase separation between multiple components ^39, 52^.

In short, we have developed a microfluidics-based chip suited for characterizing sequence-to-binodal relationships by determining precise saturation concentrations at low protein and experimental cost. This will enable further advances towards our goal of developing experiment-grounded theories of protein phase separation and will eventually enable the rational engineering of phase-separating proteins with designed solution-responsiveness.

## Materials and methods

### Device fabrication

A 4-inch silicon wafer was structured with a first SU8-3010 layer of 10 μm and a second SU8-3050 layer of 50 μm (both MicroChem), according to manufacturer instructions using a μPG101 exposure system (Heidelberg Instruments) (Fig. S1). Sylgard 184 PDMS (Dupont) was prepared as 1:10 crosslinker to base ratio and cured to a layer of 600 μm. After degassing in a rough vacuum chamber, this layer was cured at 75 °C for 1 hr. Subsequently, approximately 3 mm high and 1.5 cm by 1.5 cm wide blue casting silicon (Smooth-ON) blocks were placed onto the storage unit section of the chip and additional Sylgard was added to fill the surrounding master to a total height of 5 mm and cured at 75 °C for 1 hr. These blue silicon squares were then cut out with a tweezer. Alternatively, adhesive tape was used to shield their sides from reacting with the second PDMS cast. After punching a fluid port through the 5 mm thick Sylgard using 0.75 mm diameter biopsy punches (World Precision Instruments) the PDMS chip was bonded onto glass substrates using an 0.3 mbar, 50% power, 15 second O_2_-plasma exposure using a Diener Zepto plasma machine (Diener Electronics). The blue casting silicon patches where removed directly after bonding and channels were immediately purged with a 1:20 dilution of Cytop 809M in CTsolv180E (both Asahi Glass). The chip was then baked at 180°C for 1 hr to allow for covalent bonds to form between the plasma activated channel surfaces and the Cytop polymer. After cooling down to room temperature, chips were stored until use.

### Device loading

A detailed loading instruction is provided in Fig. 2. A ∼4 cm section of 0.3/0.75 mm ID/OD PTFE tubing (Novodirekt GmbH) was inserted into all channel outlet punch holes. Outlet tubings where terminated into a P-881 fitting with a dead-end plug (IDEX). All inlets where prepared with an identical tubing segment capped with a PDMS cube of 5 mm edge length with a 0.75 mm punch-through hole. An identical tubing segment capped by a PDMS cube was prepared as the injection loop receiver.

Channels and connected inlet and outlet tubing where primed by flushing 25 μl droplet oil (BioRad) using a 25 μl direct displacement pipette (Gilson) from the outlet through the complete device until no air bubbles remained trapped in the complete fluidic circuit. The outlet was then sealed by inserting the P-881 dead-end plug. Then sample was aspirated into a 2 μl manual air displacement pipette (Gilson) and the injection loop was connected to the 2 μl pipette tip (Fig. 2A) to inject the sample. The tip was then ejected from the pipette and the injection loop was connected to the inlet tube. After removal of the tip, a hydrostatic pressure feed HFE-7500 (3M) reservoir was connected to the injection loop. After positioning the chip on the microscope, the P-881 dead-end plug was opened slightly to allow for the hydrostatic pressure-driven flow to load the wells. After all wells had filled, the dead-end plugs where closed again.

### Constructs

Amino acid sequence of the three proteins used in this study are listed below. The coding sequences for A1-LCD and A1-LCD+12D+7R (with 12 additional aspartates and 7 arginines) including an N-terminal ENLYFQGS TEV protease cleavage site were synthesized with attB sequences at the 5’ and 3’ ends for Gateway cloning. A1-LCD and A1-LCD+12D+7R genes were cloned into pDEST17 vector (Invitrogen) which includes an N-terminal 6xHis-tag coding sequence. In the expressed protein, the N-terminal 6xHis-tag was cleaved using the TEV protease cleavage site, leaving only an additional GS sequence (underlined). BSA (UniProt ID: P02769) was purchased from Sigma.

A1-LCD:

<u>GS</u>MASASSSQRGRSGSGNFGGGRGGGFGGNDNFGRGGNFSGRGGFGGSRGG GGYGGSGDGYNGFGNDGSNFGGGGSYNDFGNYNNQSSNFGPMKGGNFGGRSSGPYGGGGQYFAKPRNQGGYGGSSSSSSYGSGRRF

A1-LCD^+12D+7R^:

<u>GS</u>MASADSSQRDRDDRGNFGDGRGGGFGGNDNFGRGGNFSDRGGFGGSRGDG RYGGDGDRYNGFGNDGRNFGGGGSYNDFGNYNNQSSNFDPMKGGNFRDRSSGPYDRGGQYFAKPRNQGGYGGSSSSRSYGSDRRF

### Sample preparation

Polyethylenglycol (PEG) 10,000 was purchased from Roth and BSA from Sigma. BSA was dissolved in ddH_2_O to obtain a stock solution of 100 mg/ml. All pairwise PEG and ammonium sulfate (AS) (% w/w) concentrations were prepared off-chip. A1-LCD was expressed and purified as previously described ^23^. The buffer exchange into native-like conditions of A1-LCD and A1-LCD^+12D+7R^ was achieved in two steps. The protein in 6 M GdmHCl, 20 mM MES pH 5.5 storage buffer was first exchanged into 1 M MES pH 5.5 by multiple dilution and concentration steps using a 3000 MWCO Amicon centrifugal filter. The protein was then dialyzed overnight against 20 mM HEPES pH 7.0 at room temperature. The protein was filtered through a 0.22 μm Millex-GV filter (Merck) to remove potential aggregates, which might have formed during dialysis. The pH of the buffer was adjusted using dilute ammonium hydroxide to prevent the introduction of excess salt into the sample and prevent phase separation. Protein concentrations were determined by absorbance at 280 nm on a NanoDrop spectrophotometer using extinction coefficients of 43824 M^-1^ cm^-1^ for BSA, 11920 M^-1^ cm^-1^ for A1-LCD and A1-LCD^+12D+7R^.

### Device N-terminal fluorescent labeling of primary amines on proteins

A1-LCD was fluorescently labeled on the N-terminus using Alexa Fluor® 488 NHS ester fluorescent dye (ThermoFisher) to label the primary amines of proteins. The protein in 6 M GdmHCl, 20 mM MES pH 5.5 storage buffer was exchanged into 20 mM HEPES pH 7.0 as described above. The protein solution was mixed with the dye using a 1 to 3 molar ratio, the mixture was incubated in the dark for 1h at room temperature before removing the excess dye via dialysis against 20 mM HEPES pH 7.0.

### Construction of binodals from UV absorbance and light microscopy

We mapped binodals in test tubes to validate the saturation concentrations measured on chip. LLPS was induced by adding NaCl to different final concentrations to the protein in storage buffer 20 mM HEPES pH 7.0 without excess salt. The resulting protein solution was incubated at 20°C for 20 min followed by centrifugation at 15,000 rpm for 5 min in a temperature-controlled centrifuge. *C*_*sat*_ was determined by removing the dilute phase and determining its absorbance at 280 nm on a NanoDrop (Thermo) UV-Vis spectrophotometer. The error bars in the figures are the standard deviation of three replicate measurements incubated at the same temperature. 2 μL of the dilute phase were diluted with 2 μL buffer before loading onto the chip. All mixing chambers were monitored in real time using a Zeiss LSM 780 or LSM 800 confocal microscope at room temperature. Sample images were taken every three or five minutes using a 10x or 20x objective. Continuous evaporation of water over time, while the temperature was kept constant, resulted in increased protein and solute concentrations, eventually resulting in LLPS for some conditions. The chamber is a cylinder and the volume (*V*) of a cylinder is expressed by *V = πr*^*2*^*h*, where *r* is the radius and *h* is the height of the cylinder. We know the protein concentration inside the chamber at time point t_0_ and its volume (Fig. 3). When water evaporates, the protein solution becomes more concentrated and *r* smaller. The water is replaced by oil in the chamber. The chambers are designed so that only the radius of the protein solution changes while the height remains constant. This allows the calculation of the protein and NaCl concentration when phase separation occurs (at *t*_sat_) from the droplet area. Three droplets per condition were analyzed using the open source image processing package Fiji ^53^ based on ImageJ and the mean was reported as *c*_sat_. The error of *c*_sat_ was calculated as the standard deviation of the mean.

## Acknowledgements

We gratefully acknowledge Petra Schwille for generously providing access to the microfabrication and microscopy lab resources. M.H. acknowledges support from the Heinrich Herz foundation through an Add-On Fellowship and project number “402723784” funded by DFG, German Research Foundation. T.M. acknowledges funding by NIH grant R01GM112846, by the St. Jude Children’s Research Hospital Research Collaborative on Membrane-less Organelles in Health and Disease, and by the American Lebanese Syrian Associated Charities. The content is solely the responsibility of the authors and does not necessarily represent the official views of the National Institutes of Health.

