## supplemental text for "Microfluidic characterization of macromolecular liquid-liquid phase separation"

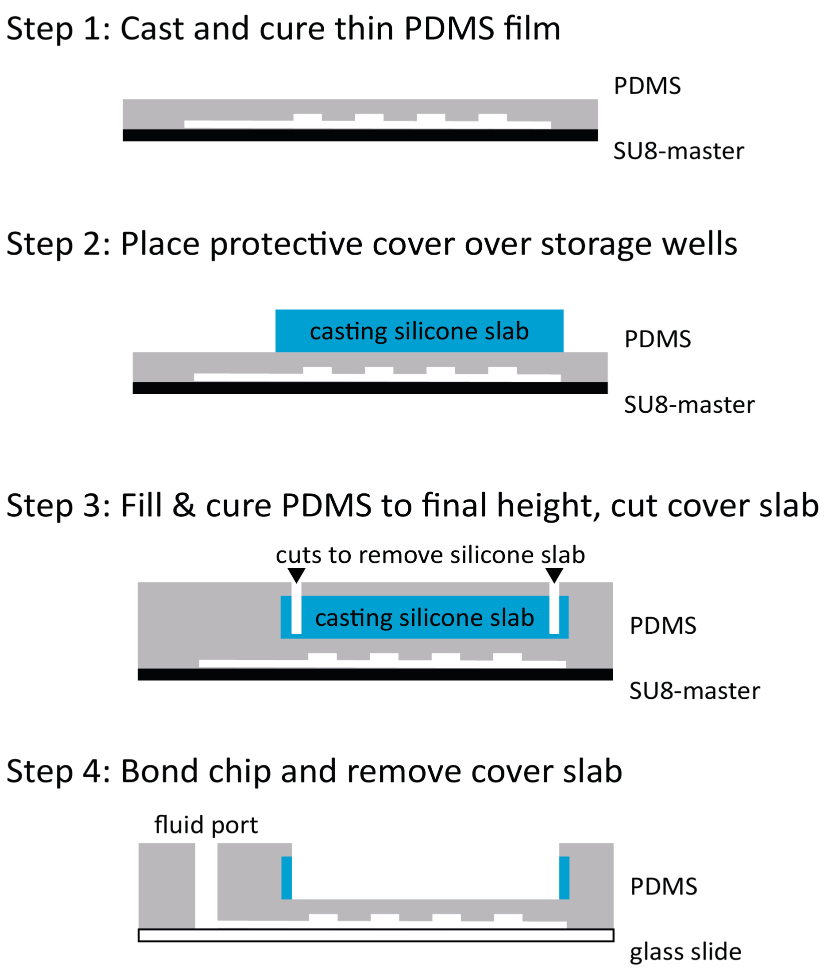


**Fig. S1: Fabrication schematic**. **Step 1**: Pour a 600 μm PDMS layer, degas for 15 min and cure at 75 °C for 1 hr. **Step 2**: Place an approximately 3 mm high and 1.5 cm by 1.5 cm wide blue casting silicon blocks onto the storage unit section of the chip. Adhesive tape may be used to mask the sides from reacting with the subsequent PDMS cast to facilitate later removal of the slab. **Step 3**: Add PDMS to a total height of 5 mm and cure at 75 °C for 1 hr. Cut the blue silicon squares with a scalpel. **Step 4**: Punch fluid ports and bond the chip onto a glass substrate. Remove blue casting silicon patches directly after bonding and purge channels immediately with a 1:20 dilution of Cytop 809M in CTsolv180E. Bake chip at 180 °C for 1 hr to allow for covalent bonds to form between the plasma activated channel surfaces and the Cytop polymer. Allow chips to cool down to room temperature and store until use.

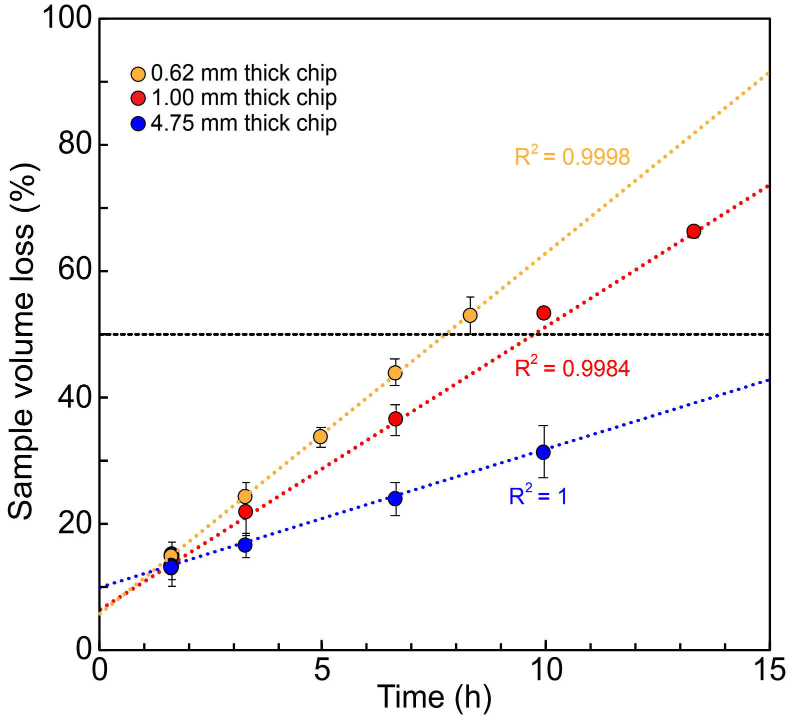


**Fig. S2: The sample volume loss over time depends on chip thickness.** The sample volume loss of 20 mM HEPES, 150 mM NaCl pH 7.0 for three different chip thicknesses was analyzed. The horizontal line indicates 50% sample volume loss. Solvent evaporates faster from a thinner chip. The regression line does not go through zero because there was a lack time when the sample was mounted onto the microscope and between the start of the imaging at $t_0$.

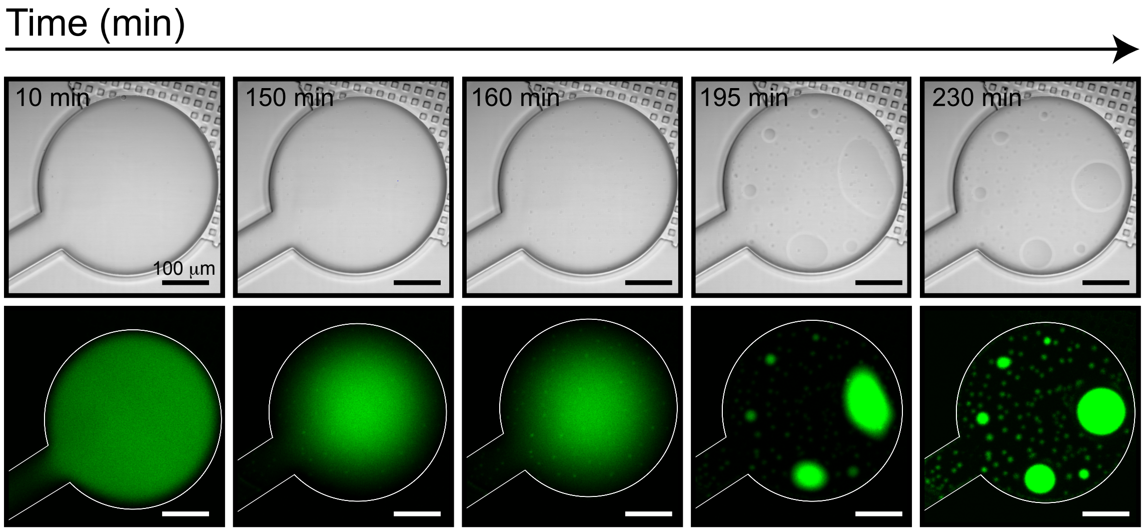


**Fig. S3:** We fluorescently labeled A1-LCD and monitored its phase separation over time to determine potential absorption of the protein to the device surface. DIC and fluorescent images are shown here. Images were taken with a LSM 800 microscope using a 20x objective. Scalebars represent 100 µm.

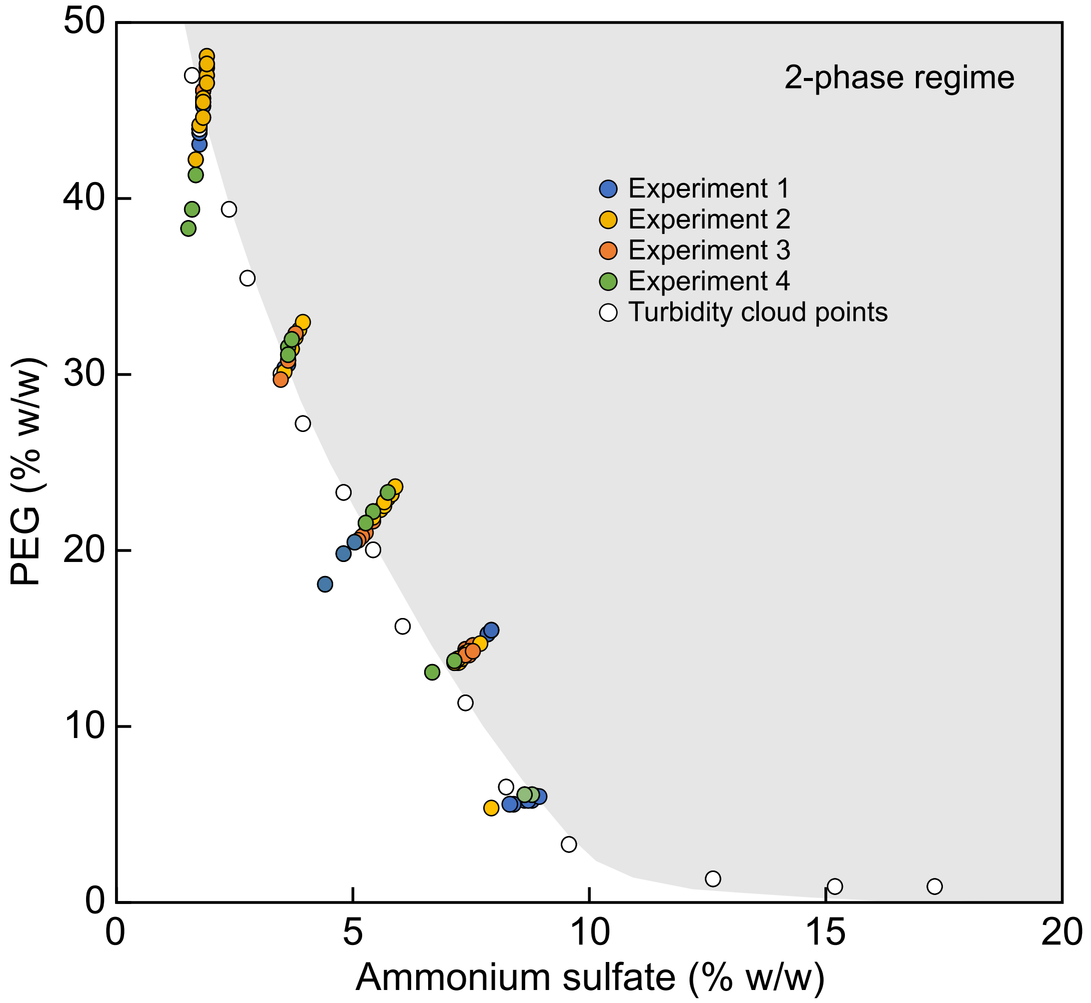


**Fig. S4: Saturation concentrations were determined from three to four independent experiments for five different pairwise PEG/ammonium sulfate combinations.** Observations from ten microfluidic chambers per condition were analyzed. A higher resolution objective (20x) was used in Experiment 4 to test whether a higher resolution allows the detection of dense phase formation earlier on. Values for $c_{sat}$ obtained using the lower *vs.* higher resolution objective were the same within error. Saturation concentrations measured on chip were compared to previously reported data points measured via turbidity cloud point assays (white circles) [45].

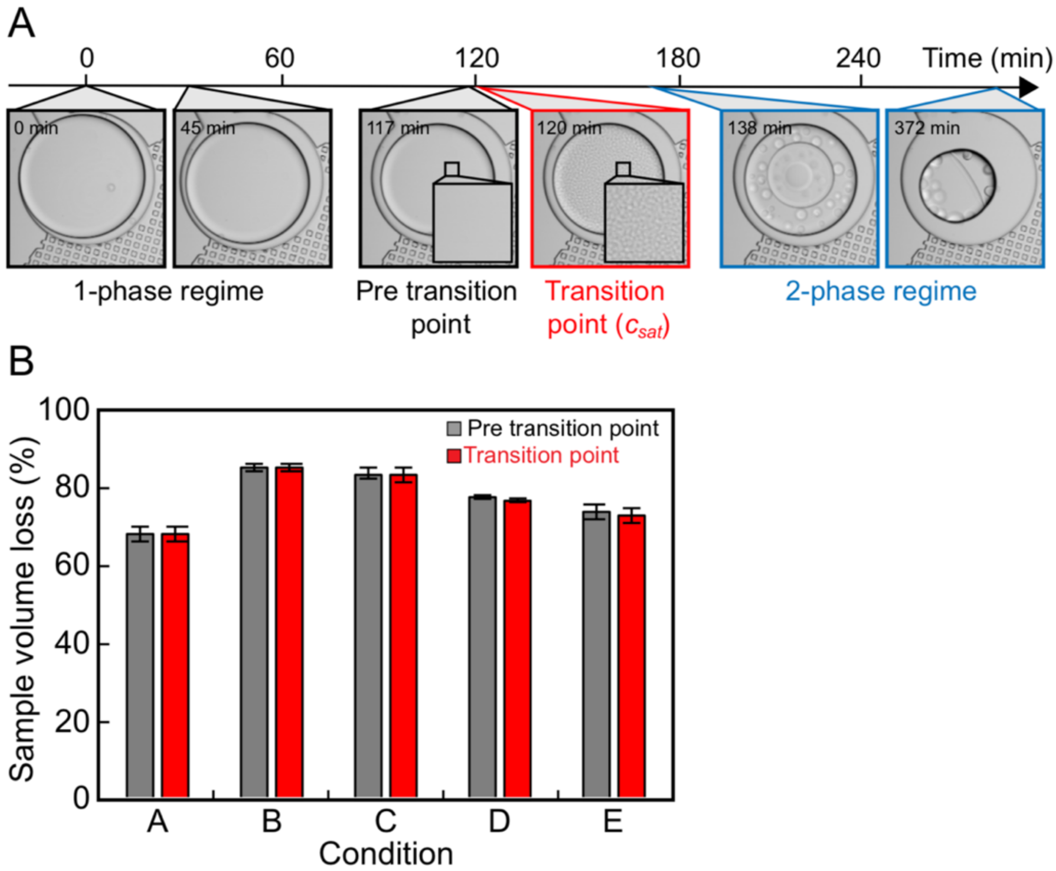


**Fig. S5: Difference in sample volume between the time point immediately before the onset of phase separation *vs*. the transition point. (A)** Snapshots of PEG/ammonium sulfate solutions taken at different time points. Images were recorded every three minutes throughout the experiment. The actual transition point may be in fact between the nominal transition point and the time point before. Therefore, we determined the difference of the sample volume between the two time points. The analysis in **(B)** shows that the size difference is less than 3% between the two time points for all five conditions measured on the chip.

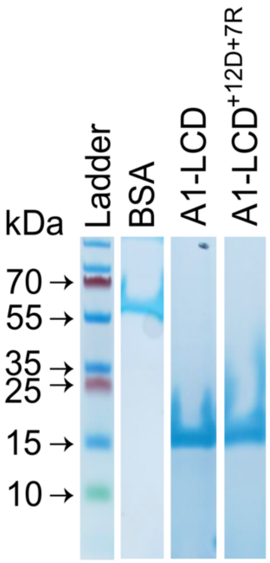


**Fig. S6:** SDS-PAGE of purified proteins used in this study.

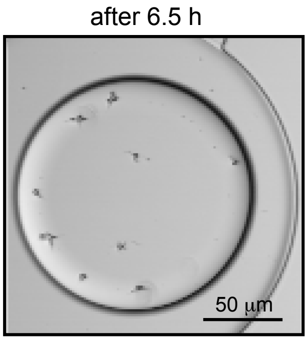


**Fig. S7:** After initial liquid-liquid phase separation, fibril-like structures were observed originating from dense phases for some A1-LCD samples.

**SI File captions**

**SI-Design file**: Design files are provided in Autodesk Autocad .DWG format. Layer 1 was fabricated from SU8-3010 to be 10 μm thick. Layer 2 was fabricated from SU8-3050 to be 50 μm for a total height of 60 μm.

**Movie S1**: Time lapse movie of the five representative PEG/ammonium sulfate filled wells shown in Figure 4 with initial concentrations A) 30/1.25, B) 23.8/2.8, C) 17.5/4.4, D) 11.3/5.9 and E) 5/7.5. Total runtime 7 hrs and 6 min. Scale bar 100 μm.

**Movie S2**: Time lapse movie of the five representative BSA/PEG wells shown in Figure 5B, E, F. Total runtime 12 hrs and 10 min. Scale bar 100 μm.

**Movie S3**: Time lapse movie of the five representative A1-LCD wells shown in Figure 5C, E, F. Total runtime 8 hrs and 15 min. Scale bar 100 μm.

**Movie S4**: Time lapse movie of the five representative A1-LCD$^{+12D+7R}$ wells shown in Figure 5D, E, F. Total runtime 13 hrs and 30 min. Scale bar 100 μm.
